# Association study of schizophrenia with variants in miR-137 binding sites

**DOI:** 10.1101/150409

**Authors:** David Curtis, Warren Emmett

**Affiliations:** UCL Genetics Institute, University College London; Centre for Psychiatry, Barts and the London School of Medicine and Dentistry; The Francis Crick Institute; Department of Molecular Neuroscience, UCL Institute of Neurology

**Keywords:** miR-137, schizophrenia, microRNA, exome, UTR, binding site

## Abstract

There is strong cumulative evidence for the involvement of miR-137 and its targets in the aetiology of schizophrenia. Here we test whether variants, especially rare variants, in miR137 binding sites are associated with schizophrenia in an exome-sequenced sample of 4225 cases and 5834 controls. A weighted burden test using 372 variants was significant at p=0.024. The sample size is too small to implicate individual variants or genes but overall this finding provides further support for the hypothesis that disruption of miR-137 binding sites can increase the risk of schizophrenia, perhaps by leading to over-expression of the target gene. These findings could be followed up by genotyping these variants in larger samples and by experimentally testing whether they do indeed effect expression. When carrying out exome sequencing it is important to include UTRs so that disruption of microRNA bindings sites can be detected.

## Introduction

MicroRNAs can bind to specific sites in the 3’ UTR of target genes and modulate their expression. As discussed recently (Olde Loohuis et al. 2017) markers for both the gene for miR-137 and for the genes it targets demonstrate association with schizophrenia (Kwon et al. 2013; Schizophrenia Working Group of the Psychiatric Genomics Consortium 2014). Experimentally, reducing or increasing miR-137 expression in rat hippocampal neurons identifies sets of regulated genes involved in neurodevelopmental processes and neuronal maturation (Olde Loohuis et al. 2017). Here, we investigate whether variants in the binding sites of miR-137 are more commonly found in exome-sequenced schizophrenia cases than controls.

## Material and methods

The data analysed consisted of whole exome sequence variants downloaded from dbGaP from a Swedish schizophrenia association study containing 4968 cases and 6245 controls (Genovese et al. 2016). As described elsewhere (Curtis 2017), the dataset was subjected to QC procedures including the removal of subjects who appeared to have a substantial Finnish component to their ancestry to leave a sample of 4225 cases and 5834 controls. As reported in the original paper, the subjects were sequenced in 12 waves. For all but the first wave the Agilent SureSelect Human All Exon v.2 Kit was used for the hybrid-capture procedure whereas for the first wave an earlier version was used. Predicted binding sites for miR-137 were obtained from <u>microRNA.org</u> (Betel et al. 2010). Excluding the Y chromosome, there were 8145 predicted binding sites, of which 1139 were covered by SureSelect, Variants in these regions were extracted and analysed by SCOREASSOC, which performs a weighted burden test such that very rare variants are weighted 10 times higher than a common variant with MAF=0.5 (Curtis 2012). Each subject receives a score consisting of the weighted sum of the variant alleles possessed by that subject at all sites and a t test is used to compare the scores between cases and controls. The predicted effects of variants were annotated using VEP (McLaren et al. 2016).

## Results

In the regions covered, 372 variants were found which passed QC procedures. On average, scores were higher for cases than controls (t=2.3, 10057 df, p=0.024), indicating that subjects with schizophrenia are more likely to have variants, especially rare variants, in miR137 binding sites than controls. The results are presented in full in Supplementary Table 1. Many variants only occurred in one or two subjects and for others there were mostly only small differences in frequencies between cases and controls so the effect could not be assigned to specific variants. However it may be worth noting the results for two variants. A variant at 10:106027165, rs7589, was heterozygous in 5 out of 4224 cases and none out of 5833 controls. It lies in the 3’ UTR of some transcripts of GSTO1. A variant at 19:58773876 was heterozygous in 17 out of 4205 cases and 13 out of 5814 controls, OR=1.8. It lies in the 3′UTR of some transcripts of ZNF544.

## Discussion

Our results provide some further support for the hypothesis that abnormalities in miR-137 functionality could be involved in the aetiology of schizophrenia. Disruption of a microRNA binding site could lead to increased gene expression, providing a mechanism for a dominant, gain of function effect. With the sample sizes used it is not possible to assign risk to individual variants and the two which we refer to are too rare to have been imputed in large GWAS samples (Schizophrenia Working Group of the Psychiatric Genomics Consortium 2014). All five subjects with the variant in the GSTO1 site have schizophrenia. The gene codes for an omega class glutathione S-transferase and a study showed that patients with schizophrenia have reduced glutathione levels in cerebrospinal fluid (Do et al. 2000) but an association study of schizophrenia with markers for GSTO1 and other glutathione related genes was negative (Matsuzawa et al. 2009). ZNF544 belongs to the C2H2-type zinc-finger family and is involved in gene transcription. In a genome-wide association study of ADHD traits an intronic SNP, rs260461, was significant at p=10^-5^ (Lasky-Su et al. 2008) and in a methylome -wide study a CpG island near ZNF544 was found to be hyper-methylated at birth in subjects with a high trajectory for ADHD symptoms (Walton et al. 2017).

In order to follow up these findings, genotyping could be carried out in the larger datasets which have been recruited for association studies. The variants could be introduced into cell cultures in order to determine whether they do indeed affect gene expression and if so in which tissues. Only a minority of binding sites were covered by the capture procedure and we recommend that exome sequencing studies should routinely include UTRs so that variants potentially affecting microRNA binding can be detected.

## Acknowledgments

The datasets used for the analysis described in this manuscript were obtained from dbGaP at http://www.ncbi.nlm.nih.gov/gap through dbGaP accession number *phs000473.v2.p2*. Samples used for data analysis were provided by the Swedish Cohort Collection supported by the NIMH grant R01 MH077139, the Sylvan C. Herman Foundation, the Stanley Medical Research Institute and The Swedish Research Council (grants 2009-4959 and 2011-4659). Support for the exome sequencing was provided by the NIMH Grand Opportunity grant RCMH089905, the Sylvan C. Herman Foundation, a grant from the Stanley Medical Research Institute and multiple gifts to the Stanley Center for Psychiatric Research at the Broad Institute of MIT and Harvard. The work of W.E. is supported by the Wellcome Trust (103760/Z/14/Z) and the Francis Crick Institute, which receives its core funding from Cancer Research UK (FC001002), the UK Medical Research Council (FC001002), and the Wellcome Trust (FC001002);

